# Assessment of extracellular vesicle protein cargo as neurodegenerative disease biomarkers

**DOI:** 10.1101/2023.11.02.565137

**Authors:** Tal Gilboa, Dmitry Ter-Ovanesyan, Shih-Chin Wang, Sara Whiteman, George T. Kannarkat, George M. Church, Alice S. Chen-Plotkin, David R Walt

**Affiliations:** Wyss Institute for Biologically Inspired Engineering, Boston, MA, USA; Department of Pathology, Brigham and Women’s Hospital, Boston, MA, USA; Harvard Medical School, Boston, MA, USA; Department of Neurology, Perelman School of Medicine at the University of Pennsylvania, Philadelphia, PA, USA

## Abstract

Extracellular vesicles (EVs) are released by all cells and hold great promise as a class of biomarkers. As EVs represent a way of capturing molecular information about the proteins inside of cells, EVs from biofluids could be used to better understand and diagnose disease from difficult to access organs such as the brain. This promise has led to increased interest in measuring EV proteins from both total EVs as well as brain-derived EVs isolated from the blood. However, the measurement of cargo proteins in EVs has been challenging because EVs are present at low levels and EV isolation methods are imperfect at separating EVs from free proteins. Thus, it is difficult to know whether a protein measured after EV isolation is truly inside EVs. In this study, we developed methods to measure whether a protein is inside EVs and quantify the ratio of a protein in EVs relative to total plasma. To achieve this, we combined a high-yield size exclusion chromatography (SEC) protocol with an optimized protease protection assay and Single Molecule Array (Simoa) digital ELISA assays for ultrasensitive measurement of proteins inside EVs. We applied these methods to analyze key proteins involved in neurodegenerative diseases: α-synuclein, Tau, Aβ40, and Aβ42. We found that α-synuclein and Tau are present in plasma EVs at a small fraction of the levels in total plasma, whereas Aβ40 and Aβ42 are undetectable in plasma EVs. This work provides a framework for determining the levels of proteins in EVs and represents an important step in the development of EV diagnostics for diseases of the brain, as well as other organs.

## Introduction

Extracellular vesicles (EVs) are released into circulation and contain cargo molecules from their donor cells (1). Since all cells release EVs, analyzing the contents of EVs from biofluids such as plasma provides an avenue to determine the state of inaccessible organs and diagnose pathology at the molecular level (2). Additionally, as EVs contain proteins from the donor cell, a potential advantage of using EVs is that cargo molecules can be linked to the cell of origin (3).

Since it is very high risk to biopsy the brain in a clinical setting, neuron-specific EVs are particularly promising biomarkers for neurodegenerative diseases (4). Analyzing proteins involved in neurodegenerative disease inside of neuron-specific EVs isolated from plasma could enable diagnostics for early detection or progression monitoring of diseases such as Parkinson’s Disease (PD) and Alzheimer’s Disease (AD) (5, 6). One such protein of interest is α-synuclein, which when misfolded, aggregates, and spreads along neuronal networks is associated with onset and progression of PD (7). α-synuclein is highly expressed in the brain but also in red blood cells (8). Thus, the majority of α-synuclein measurable in plasma is thought to originate from red blood cells. Measuring α-synuclein from neuron-specific EVs would more likely mirror the pathological form of α-synuclein in the brain (9). α-synuclein phosphorylated at the Ser129 residue (pSer129), for example, is considered to be a pathogenic form of α-synuclein associated with PD (10, 11). Similarly, in AD, proteins such as Tau, Aβ40, and Aβ42 are associated with disease progression in the brain. Cerebrospinal fluid (CSF) measures of phosphorylated and unphosphorylated forms of Tau, as well as Aβ40, and Aβ42, are already used diagnostically for AD (12, 13), and plasma measures of these proteins and are being developed as biomarkers (14).

A large number of studies have reported the measurement of various proteins after immuno-isolation of neuronal EVs from plasma (4-6), but several technical challenges complicate the interpretation of these results. In particular, a neuronal EV marker for immunoisolation has not yet been established and verified. Most previous studies have used L1CAM (4, 9, 15), but we previously found that L1CAM is not associated with EVs in cerebrospinal fluid (CSF) and plasma (16). Furthermore, EV isolation, whether total or cell-type specific, often leads to co-isolation of contaminating free proteins (17). This problem is exacerbated by the fact that the levels of proteins inside EVs are often very low relative to the levels of these same proteins in their free form (18). Thus, it is unclear how much of a given protein is truly inside EVs as opposed to outside EVs (19).

In this study, we set out to determine the levels of several proteins associated with neurodegenerative diseases inside plasma EVs relative to total plasma. We used a highly optimized size exclusion chromatography (SEC) method (20, 21) to recover most EVs from plasma. We combined this high-yield EV isolation method with ultrasensitive Single Molecule Array (Simoa) immunoassays (11) to ensure we could detect the lowest possible levels of proteins inside EVs. We developed new Simoa assays for total α-synuclein and pSer129 that are much more sensitive than previously reported assays (22), and combined these assays with Simoa assays for key proteins involved in AD that have previously been reported in EVs: Tau, Aβ40, and Aβ42 (23). To ensure we were detecting EV luminal proteins, we also developed a protease protection assay to differentiate internal EV proteins from external ones. Using these methods, we were able to determine which proteins involved in neurodegenerative disease are present in total plasma EVs and provide an upper bound for the possible levels of these proteins in neuron-derived EVs.

## Results

To evaluate the contents of plasma EVs, we first selected α-synuclein as a candidate EV cargo protein (Figure 1A). We developed a highly sensitive Simoa immunoassay for monomeric α-synuclein (SI Figure 1A). To confirm the specificity of the α-synuclein assay in biofluids, we extensively validated this assay using dilution linearity and spike and recovery experiments in plasma samples with or without fractionation (SI Figure 1B & C, SI Table 1). To measure the distribution of EV-associated and freely circulating α-synuclein, we performed SEC on plasma and used Simoa to analyze α-synuclein levels in each fraction. To precisely identify EV and soluble protein fractions (16, 21), we also used Simoa to measure the levels of a common EV marker, CD9, and an abundant protein marker, Albumin, in each SEC fraction. We found that the majority of α-synuclein was present in the later SEC fractions corresponding to free proteins, but a small amount of α-synuclein peaked in the early SEC fractions, which overlapped with the CD9 distribution (Figure 1B). This two-peak distribution of α-synuclein suggests that the first peak is EV-associated α-synuclein, while the second peak represents its freely circulating form. The second α-synuclein peak is observed later than the Albumin peak, potentially due to α-synuclein’s small size of 14 kDa (24).

**Figure 1.**
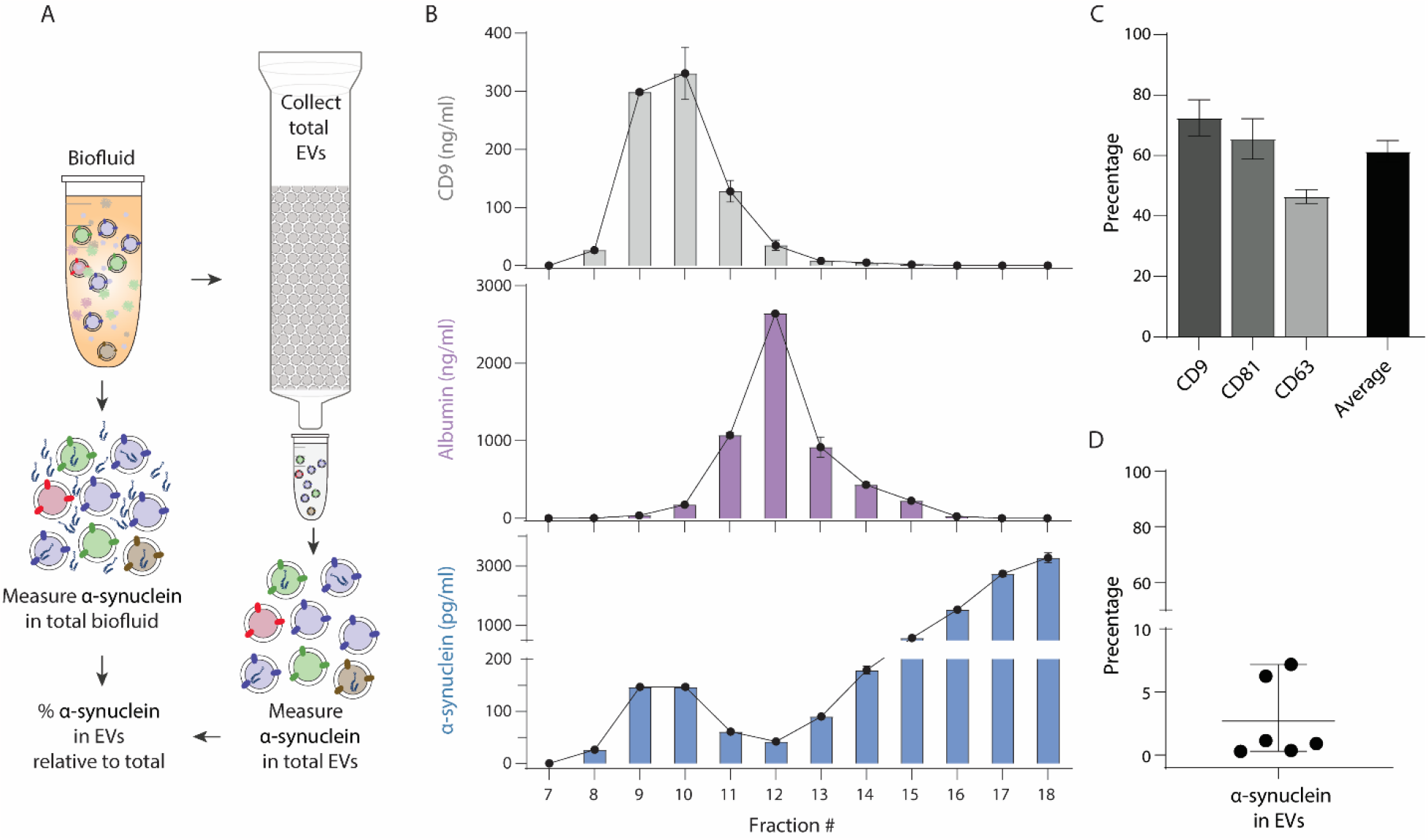
The distribution of α-synuclein in size exclusion chromatography (SEC) fractions isolated from plasma. (A) Schematic of the overall goal: comparing the level of α-synuclein in EVs relative to α-synuclein in total plasma. (B) Pooled plasma was fractionated by SEC and Simoa was used to measure levels of the EV surface marker CD9 (top), soluble protein marker Albumin (soluble protein marker, purple), and α-synuclein (blue) in each fraction. All the data are displayed as mean ± SD. (C) Levels of tetraspanins CD9, CD63, and CD81 were measured by Simoa in EVs isolated from pooled plasma by SEC (fractions 7-10) as well as in unfractionated plasma. EV recovery percentage was calculated as the average of the ratios of each tetraspanin in EVs relative to unfractionated plasma. All the data are displayed as mean ± SD of duplicate measurements. (D) Levels of α-synuclein were measured by Simoa in EVs isolated from six individual plasma samples by SEC (fractions 7-10) as well as in unfractionated plasma. The percentage of α-synuclein in the EV fractions relative to the total plasma α-synuclein was calculated for each sample.

To estimate the percentage of α-synuclein in the EVs relative to their total in plasma, we wanted to ensure we used an EV isolation method that allows for high EV recovery while depleting most free proteins. We previously used Simoa assays for EV surface proteins (the tetraspanins CD9, CD63, and CD81) to compare SEC to other EV isolation methods (20) and further optimized SEC using the Sepharose CL-6B resin to isolate EVs from plasma (21). We utilized Simoa to assess the EV recovery percentage of this isolation method by comparing CD9, CD63, and CD81 levels in SEC fractions 7-10 to those in unfractionated plasma (Figure 1C).

Having confirmed more than 50% recovery of EVs from plasma in SEC fractions 7-10, we then compared levels of α-synuclein in EVs relative to total plasma. We performed SEC on six individual plasma samples. For each sample, we used Simoa to measure α-synuclein in both SEC fractions 7-10 and unfractionated plasma (SI Figure 2). We calculated the percentage of EV-associated α-synuclein relative to the total α-synuclein in unfractionated plasma and found that this number ranges between 0.3-7% (Figure 1D).

**Figure 2.**
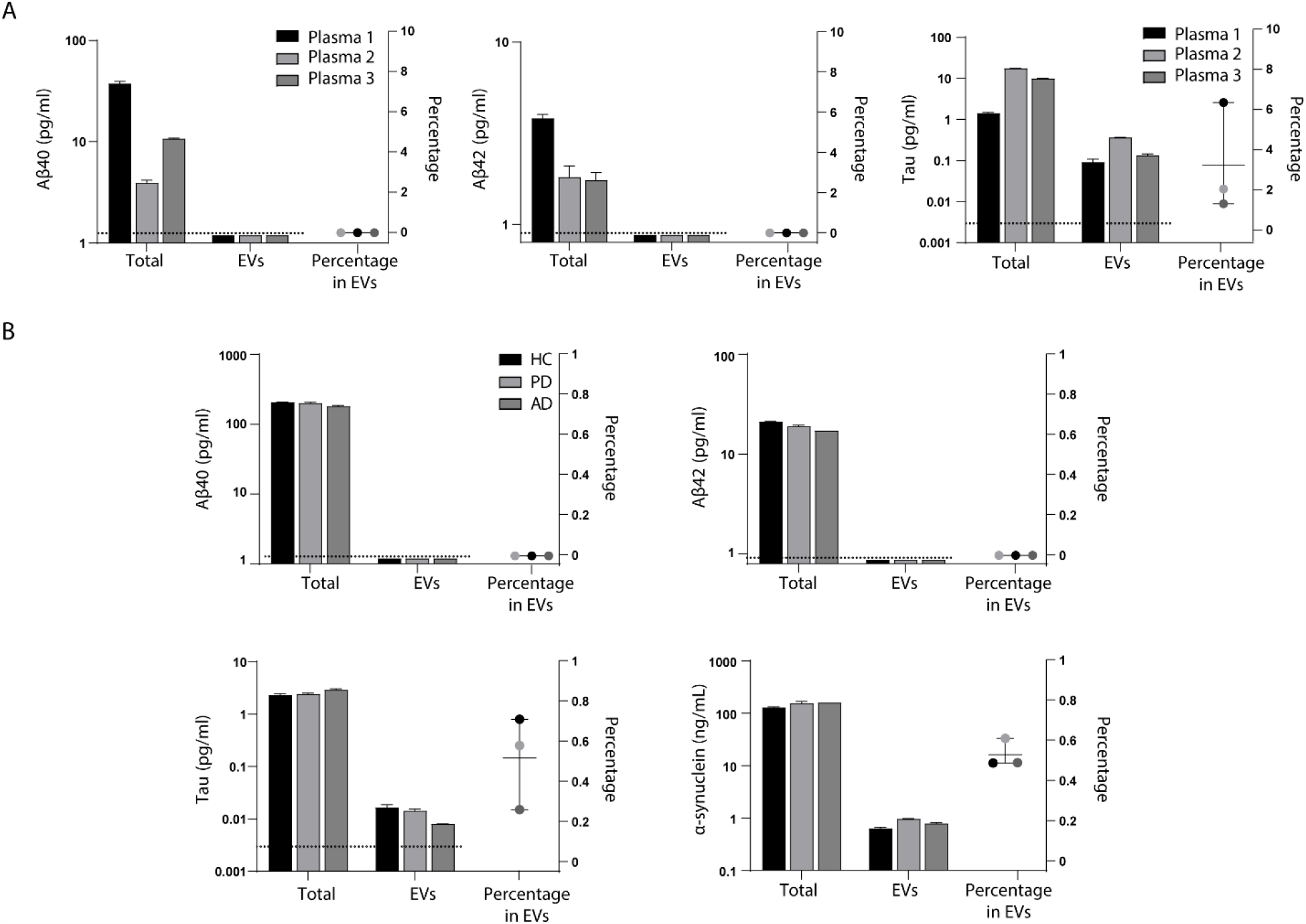
Measurement of plasma α-synuclein, Tau, Aβ40, and Aβ42 levels in healthy and disease samples and corresponding EVs isolated by SEC. Levels of (A) Aβ40, (B) Aβ42, and (C) Tau were measured by Simoa in EVs isolated by SEC (fractions 7-10) from three individual plasma samples. Levels of each protein were also measured in corresponding unfractionated plasma samples, and a percentage of each protein in EVs was calculated. All the data are displayed as mean ± SD. Dotted line indicates limit of detection for each Simoa assay. (D) Levels of Aβ40, (E) Aβ42, (F) Tau and (G) α-synuclein were measured by Simoa in EVs isolated by SEC (fractions 7-10) from pooled plasma samples of the following: healthy controls (HC), Parkinson’s Disease (PD), or Alzheimer’s Disease (AD). Levels of each protein were also measured in corresponding unfractionated plasma samples, and a percentage of each protein in EVs was calculated. All the data are displayed as mean ± SD of duplicate measurements. The dotted line indicates the limit of detection for each Simoa assay.

After measuring total α-synuclein, we applied the same approach to measure α-synuclein pSer129 levels in EVs. Expecting the levels of pSer129 to be much lower than those of total α-synuclein, we developed an ultrasensitive pSer129 Simoa assay (SI Figure 3A) and assessed dilution linearity of the assay in both EV and free protein fractions (SI Figure 3B). We also performed spike and recovery in plasma (SI Table S1). We confirmed the specificity of the assay for pSer129 by observing a loss of signal after performing phosphatase treatment to remove phosphorylation (SI Figure 3A & 3B). Using this assay, we were able to detect pSer129 in EVs isolated by SEC from four out of five individual plasma samples (SI Figure 3C). Measuring total α-synuclein in the same samples, we found the percentage of pSer129 relative to total α-synuclein in EVs to be below 0.1%, with the sample where we were unable to detect pSer129 having the lowest total α-synuclein (SI Figure 3D).

**Figure 3.**
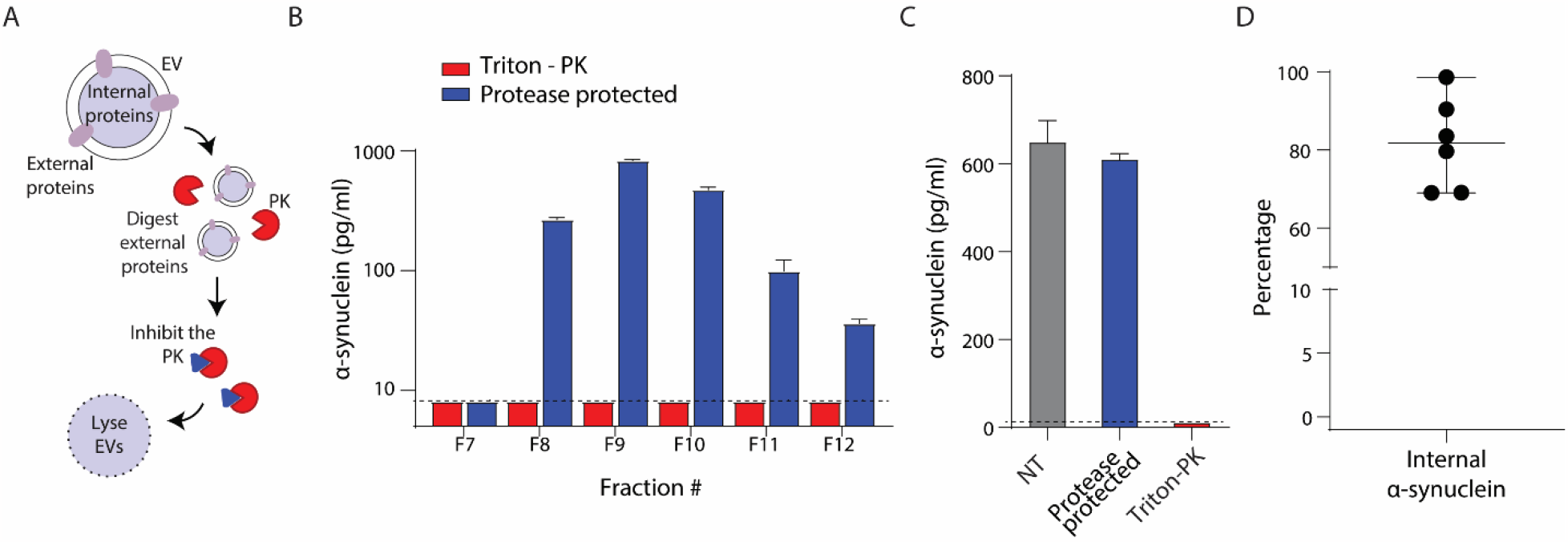
Protease protection assay for estimating the internal and external α-synuclein in plasma EVs isolated by SEC. (A) Schematic illustration of the Protease protection assay. Proteins that are not inside of EVs and protected by the lipid membrane are digested by Proteinase K (PK). Proteinase K is then inhibited and the contents of the EV are released by lysing the EVs with detergent. (B) Protease protection assay was performed on individual SEC fractions isolated from plasma. The levels of α-synuclein in each fraction were measured by Simoa after protease protection (blue bars), or with proteinase K (PK) treatment after the addition of detergent (red bars). (C) Protease protection assay was performed on pooled EV fractions (fractions 7-10; right) isolated from plasma; levels of α-synuclein were measured by Simoa in EVs isolated by SEC from pooled plasma and three conditions were compared: No treatment (NT), Protease protection assay, or PK treatment after EV lysis with TritonX-100 (Triton-PK). All the data are displayed as mean ± SD. Dotted line indicates limit of detection for α-synuclein Simoa assay. (D) The percentage of protease-protected α-synuclein (internal α-synuclein) relative to α-synuclein in untreated EVs isolated from six individual plasma samples by SEC (fractions 7-10) using the same samples as those used in Figure 1D.

Next, we applied the same methodology developed for α-synuclein to key proteins associated with AD. We used Simoa to measure Aβ40, Aβ42, and Tau levels in both EVs isolated by SEC from plasma and unfractionated plasma. While Aβ40 and Aβ42 were readily detectable in total plasma from three different individuals, Aβ40 (Figure 2A) and Aβ42 (Figure 2B) were both undetectable in the EV fractions. We were, however, able to detect Tau in EVs from these three individual plasma samples and found that Tau levels in EVs were ∼3% of Tau levels in total plasma (Figure 2C).

As the previous protein measurements of α-synuclein, Tau, Aβ40, and Aβ42 were made in EVs isolated from plasma of healthy individuals, we asked whether the levels of these proteins would be different in plasma from patients with neurodegenerative disease. We investigated this question by measuring α-synuclein, Tau, Aβ40, and Aβ42 levels in plasma EVs isolated from multiple plasma pools of PD, AD, and matched healthy controls (SI Table S2). Similar to the individual plasma samples, we could not detect Aβ40 (Figure 2D) or Aβ42 (Figure 2E) in the EV fractions of any pooled samples but were able to detect both Tau (Figure 2F) and α-synuclein (Figure 2G) in these EV fractions.

Having detected α-synuclein and Tau in the EV-containing early SEC fractions, we sought to confirm that these proteins were truly inside EVs. One general challenge in studying the contents of EVs is the inability to distinguish luminal EV cargo from free protein contaminants that are co-purified with EVs during the isolation procedure (17, 18). Such impurities could lead to misinterpretation of EV-associated biomarker discovery results and are likely present to some degree in all EV isolation techniques, including SEC (25). To overcome this challenge, we optimized a protease protection assay that degrades proteins that are not protected by the EV lipid membrane (17) (SI figure 4). In this protocol, isolated EVs are first incubated with Proteinase K (PK), followed by PK inhibition using phenylmethylsulfonyl fluoride (PMSF). Then, EVs are permeabilized with Triton X-100 detergent, releasing protected luminal proteins for downstream analysis (Figure 3A). To ensure complete inactivation of PK by PMSF, we measured α-synuclein levels using three conditions applied to either purified α-synuclein protein or isolated EVs from plasma permeabilized with detergent: PK, PK inactivated with PMSF, or no treatment (SI Figure 4B & C). We also validated the protease protection assay using additional proteins(SI Figure 4D)

**Figure 4.**
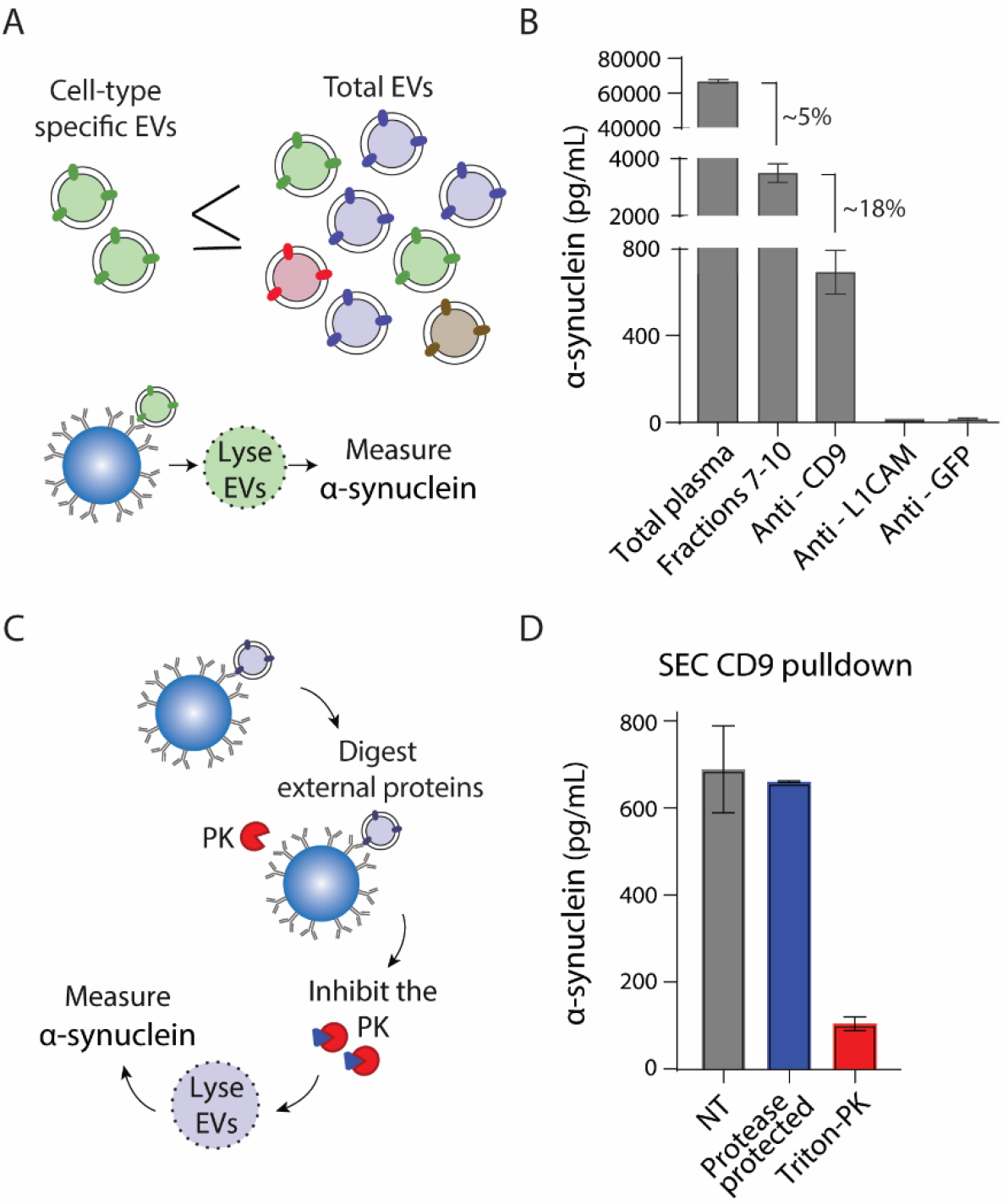
Measurement of α-synuclein in immuno-isolated subset of plasma EVs. (A) Schematic illustrating that cargo protein (α-synuclein) in an immuno-isolated subset of EVs (such as cell type specific EVs), must be less than or equal to the level of that cargo protein in total EVs. (B) Measurement of α-synuclein by Simoa in: total plasma, EVs isolated from plasma by SEC (fractions 7-10), EVs isolated by SEC from plasma (fraction 7-10) and then immuno-isolated using anti-CD9 antibody, anti-L1CAM antibody, or anti-GFP antibody. All the data are displayed as mean ± SD. (C) Schematic illustrating Protease protection assay after EV immuno-isolation to ensure cargo protein is inside EVs. (D) Levels of α-synuclein were measured by Simoa in EVs immuno-isolated using anti-CD9 antibody after SEC (collecting fractions 7-10) from pooled plasma and three conditions were compared: No treatment (NT), Protease protection assay, or PK treatment after EV lysis with Triton X-100 (Triton-PK). All the data are displayed as mean ± SD of duplicate measurements.

We applied the optimized protease protection assay to individual SEC fractions to assess whether the α-synuclein detected in the early SEC fractions is truly contained within the EVs. Measuring α-synuclein by Simoa, we observed a similar EV-associated peaking pattern in fractions 7-12 after the protease protection as we previously observed without PK treatment (Figure 3B). We did not detect α-synuclein when adding detergent to EVs before the same PK treatment, suggesting α-synuclein is protected by the EV lipid membrane. To more precisely quantify the luminal EV-associated α-synuclein, we collected SEC fractions 7-10 from pooled plasma and used Simoa to measure α-synuclein in the following conditions: no treatment (total levels of EV-associated α-synuclein), the Triton-PK condition (any potential background α-synuclein levels after digestion of both internal and external proteins in EV fractions), and the PK-PMSF-Triton protease protection condition (levels of α-synuclein inside the EVs) (Figure 3C). Performing this experiment with EVs isolated from six individual plasma samples, we found that ∼80% of EV-associated α-synuclein was protease protected (Figure 3D).

Having validated the protease protection assay and confirmed that α-synuclein in the EV fractions is mostly inside EVs, we applied the same approach to investigate Tau in plasma EVs. We used Simoa to measure Tau after performing the protease protection assay on SEC fractions 7-10 and found that ∼63% of Tau in these SEC fractions is inside EVs (SI figure 5).

Measuring the level of a cargo protein such as α-synuclein or Tau in total EVs provides an upper bound for the protein levels that can be expected in a subset of EVs. This subset of EVs can be immuno-isolated using antibodies against a cell-type specific (neuronal, etc.) protein marker (Figure 4A). To investigate protein cargo in immuno-isolated EVs, we measured α-synuclein after immuno-isolation. First, we isolated total EVs from plasma by SEC (fractions 7-10). Then we performed immuno-isolation using beads coated with antibodies against CD9, L1CAM or GFP. We lysed EVs with detergent and measured α-synuclein by Simoa. CD9, considered a general EV marker, is expressed in many cell types (26), including red blood cells (27, 28), which produce high levels of α-synuclein (8). L1CAM is highly expressed in neurons and has been extensively used for EV immuno-isolation, although we found it exists predominantly in plasma as a free protein (16). We used an anti-GFP antibody as a control since we expect some level of non-specific binding to antibody-coated beads regardless of which antibody is used. In the same experiment, as before, we also measured α-synuclein in total plasma and EVs isolated by SEC. We found that the level of α-synuclein after CD9 immuno-isolation is 18% of total EV-associated α-synuclein. The levels of α-synuclein after L1CAM or GFP immuno-isolation were both similarly low and just slightly above the limit of detection (Figure 4B).

We applied our protease protection assay to confirm that the cargo protein we detect after EV immuno-isolation is inside EVs. By treating immuno-isolated EVs with our optimized protease assay, we could eliminate α-synuclein or any other protein that may be non-specifically bound to antibody-coated beads (Figure 4C). When measuring α-synuclein after CD9 immuno-isolation from SEC fractions 7-10, we found that α-synuclein in the protease-protected condition is present at the same level as in the untreated condition (Figure 4D). Thus, we confirmed that we could detect α-synuclein inside EVs after immuno-isolation. These experiments provide a framework for measuring α-synuclein or other cargo proteins in EV subpopulations, which in the future could include cell-type-specific EVs.

## Discussion

Measuring proteins in plasma EVs relative to total plasma may have unique advantages for biomarker discovery and molecular diagnostics. First, EVs may more accurately reflect the protein state (for example, post-translational modifications) of their cell of origin (29). Second, isolating EVs from plasma depletes Albumin and other highly abundant proteins that may interfere with some types of measurement (such as proteomics) (25). Third, a cargo protein in EVs may be tied to its cell of origin by surface proteins present on the EVs (3-5). However, a crucial question in the EV biomarker field is how much cargo proteins plasma EVs contain and how this amount compares to the protein level in total plasma (18, 19, 25, 30, 31). This fundamental question was previously hampered by several challenges in the EV field: lack of consensus regarding yield and purity of different EV isolation methods (18, 29, 30), inability to distinguish whether proteins detected after EV isolation are truly inside EVs (as opposed to non-specifically bound to the outside) (17, 18, 25), and low levels of EV cargo proteins that are challenging to detect with conventional techniques (3, 31).

In this study, we brought together multiple technological advances to quantify the fraction of proteins in EVs relative to plasma. Our work demonstrates a new approach to studying the protein cargo in EVs purified from plasma. We used α-synuclein, the key protein in PD, as well as Tau, Aβ40, and Aβ42, key proteins associated with AD, to demonstrate the approach. First, we used an SEC-based EV isolation method that we have extensively optimized to recover most EVs from plasma, while efficiently depleting soluble proteins (20, 21). Second, we developed and validated a protease protection assay to ensure that we detected the cargo proteins inside EVs after SEC or immuno-isolation. Third, we used Simoa immunoassays that are highly sensitive and have a wide dynamic range (32).

To investigate α-synuclein, we developed an ultrasensitive Simoa assay to measure α-synuclein levels in plasma fractionated by SEC. Our results revealed the presence of a population of EV-associated α-synuclein as evidenced by a small but distinct peak of α-synuclein in the early SEC fractions, separated from the main peak of free α-synuclein in the later SEC fractions. Although α-synuclein and other proteins involved in neurodegenerative diseases have previously been detected in both total EVs (33) and EVs after immuno-isolation with anti-L1CAM antibody (9), it remains unclear whether protein measurements in these studies were from proteins that were truly inside EVs or non-specifically bound to the outside of EVs. We have previously reported that L1CAM in plasma is a soluble protein and not associated with EVs (16), raising doubts about whether proteins such as α-synuclein detected after L1CAM immuno-isolation in plasma are inside EVs (15). Indeed, in the present study, we did not detect α-synuclein in EV fractions subjected to L1CAM pulldown, despite using high-sensitivity Simoa assays. To assess whether α-synuclein in total EVs isolated by SEC was truly inside the vesicles, we used a protease protection assay to degrade proteins not within the lumen of the vesicles. This assay confirmed that α-synuclein in early SEC fractions is truly inside EVs. Additionally, we developed a Simoa assay for α-synuclein pSer129, and, despite the low levels of phosphorylation, were able to detect pSer129 in plasma EVs.

We extended the same approach used for α-synuclein to investigate other proteins involved in neurodegeneration. Using SEC and protease protection, we showed that there is a population of Tau present inside EVs. Unlike α-synuclein and Tau, we could not detect Aβ40 or Aβ42 in plasma EVs. Aβ40 and/or Aβ42 have been detected after L1CAM immuno-isolation in plasma (34-42), but as Aβ40 and Aβ42 are extracellular (43), it is unlikely they are inside EVs. Although these proteins may associate with the outer surface of EVs as the EVs are released from neurons, the absence of Aβ in our SEC EV fractions does not support this hypothesis. Thus, a more likely explanation is that freely circulating, non-EV-associated Aβ sticks to anti-L1CAM antibody-coated beads. This binding may occur through a non-specific interaction with the beads. Alternatively, Aβ may be binding to (immunocaptured) soluble L1CAM, which has been previously reported (44).

Although α-synuclein and Tau are present in EVs, the levels of each in EVs relative to total plasma are low. We found the percentage of α-synuclein and Tau in EVs is two and three percent, respectively, of the corresponding protein levels in plasma. As we have demonstrated that our SEC method recovers more than 50% of EVs in plasma (and most α-synuclein and Tau in the EV fractions is protected from protease), we can use these α-synuclein and Tau measurements to set an upper limit on how much of each protein can be present in cell-type specific EVs. Although we do not know what fraction of total EVs in plasma comes from a specific cell type, we expect the fraction of neuron-derived EVs to be low. Thus, having a limit of detection for our α-synuclein Simoa assay that is over a hundred times lower than what is detected in plasma EVs maximizes the chances that α-synuclein can be detected in neuron EVs. To demonstrate that we can detect α-synuclein in a subset of EVs, we also measured α-synuclein after CD9 immuno-isolation and performed protease protection to confirm that α-synuclein was inside. We could not detect α-synuclein above background levels after L1CAM immuno-isolation, which is what would be expected if L1CAM is not present on EVs. Although CD9 is widely expressed across cell types, this experimental approach can be applied in the future to immuno-isolation of neuron-specific EVs using new markers.

The isolation of cell-type specific EVs and the measurement of cargo molecules inside these EVs holds great promise for molecular diagnostics (4, 5). Before this goal can be successfully achieved, however, it is important to establish methods that ensure that a given EV cargo molecule is truly inside EVs and not non-specifically bound. In this work, we have established an experimental framework for measuring the proportion of cargo molecules in plasma EVs relative to the total biofluid and confirmed that these cargo molecules are truly inside EVs. This work represents an important step toward developing EV-based diagnostics to non-invasively measure cargo molecules in a cell type-specific fashion from the brain and other organs.

## Methods

### Human samples

Pooled and individual healthy human plasma samples were obtained from BioIVT. Plasma pools from healthy controls, PD, and AD were collected through the MIND Initiative at the Perelman School of Medicine, University of Pennsylvania (45). Plasma pools were generated by thawing individual samples on ice, combining plasma, gentle end-over-end inversion for 30 minutes at 4°C and subsequent aliquoting and storage at -80°C. Plasma was thawed on ice, centrifuged at 2,000 x g for 10 minutes at 4°C, and the supernatant was collected. The supernatant was then filtered through a 0.45 μm Costar Spin-X centrifuge tube (Millipore Sigma) and centrifuged for an additional 10 minutes at 2,000 x g at 4°C.

### Simoa assays

Simoa assays for CD9 and Albumin were conducted as previously described (21). For the total α-synuclein assay, clone 4B12 (807801, BioLegend) was used as the capture antibody and clone EPR20535 (ab225866, Abcam) was used as the detector antibody. The protein standard for the assay was TP310606 (Origene). For the pSer129 α-synuclein assay, clone D1R1R (23706, Cell Signaling Technology) was used as the capture antibody, and clone 4B12 (807801, BioLegend) as the detector antibody. The pSer129 protein standard was obtained from Proteos (RP-004). Both assays underwent validation as described in the supporting information. CD9, α-synuclein and pSer129 were measured with a two-step assay, while Albumin was measured with a three-step assay. Neurology 3-Plex A (N3PA) Assay kit (Quanterix) was used to measure Tau, Aβ40 and Aβ42. For measuring proteins in SEC fractions, the fractions were diluted 4X, with an additional 4X dilution for CD9. For measuring protein levels in total plasma, α-synuclein and CD9 were 100X diluted, and pSer129 was 20X diluted. All samples were measured in duplicate using the HD-X analyzer (Quanterix). Average Enzyme per Bead (AEB) values were calculated by the HD-X software.

### Protein phosphatase treatment

The pSer129 α-synuclein assay was validated using a phosphatase treatment. Purified pSer129 proteins or diluted soluble protein fractions isolated via size-exclusion chromatography (SEC) were subjected to protein phosphatase treatment. In a 50 μl phosphatase reaction, 5 μl of 10X PMP buffer, 5 μl of 10mM MnCl2, and 1 μl of Lambda protein phosphatase (P0753L, New England BioLabs) were incubated with the proteins at 30°C for 30 minutes.

### Protease protection assay

For the protease protection assay on EVs isolated by SEC, the biofluid was divided into three groups: no treatment (NT), protease-protected, and PK-triton. In the protease-protected and PK-triton groups, Proteinase K (PK) (E00491, Thermo Fisher Scientific) was added to the biofluid at a final concentration of 0.2 mg/mL. For the NT group, an equivalent volume of PBS was added instead of PK. Additionally, 0.5% Triton X-100 (93443, Sigma-Aldrich) was added to the PK-triton group to burst the EVs. All three groups were incubated at 37°C for 30 minutes. After incubation, 0.1M Phenylmethylsulfonyl fluoride (PMSF) (10837091001, Roche) dissolved in isopropanol was added to each group. Subsequently, samples were shaken at room temperature at 300 rpm for one hour. The NT and protease-protected groups were then supplemented with 0.5% Triton X-100. For the protease protection assay on the beads, beads were conjugated, washed and incubated with biofluid as before. The beads were resuspended in 600uL of isolation buffer then split into 3 groups (NT, PK, PK-triton). PK and PK-triton were added directly to the beads and incubated at 37°C for 30 minutes. PMSF was added to all groups then incubated while shaking at RT. 0.5% Triton X-100 was added directly to the beads. The beads were incubated for 10 minutes then the supernatant was removed and collected for further analysis.

### EV isolation

Extracellular vesicles (EVs) were isolated from plasma through size-exclusion chromatography (SEC) (21). Sepharose CL-6B resin was washed three times with an equal volume of PBS in a glass container. Subsequently, the resin was packed into 10 mL Econo-Pac Chromatography columns (Bio-Rad), and a frit was inserted atop the resin to prepare the column. The column underwent two rounds of washing with 5 mL of PBS each time to remove leftover ethanol.

Once the PBS from the wash had completely passed through the column, 1 mL of plasma was introduced onto the column. Upon the plasma sample fully entering the column, PBS was added to the column at 0.5 mL increments for collection of individual fractions. In cases where the goal was to isolate total EVs, 2 mL of PBS was introduced, and Fractions 7-10 were collected and pooled.

### EV Immuno-Isolation

For EV immuno-isolation from plasma, 1X PBS pH 7.4 was utilized as the isolation buffer. Dynabeads Goat Anti-Mouse IgG beads (Thermo Fisher Scientific) were placed into a 2 mL tube and magnetically separated. The beads were washed once with 1 mL of isolation buffer and then incubated overnight at 4°C with 10 μg of either anti-CD9 (clone CBL162, Millipore), anti-L1CAM (clone 5G3, BD) or anti-GFP (clone 9F9.F9, Rockland) antibody in 0.5 mL of isolation buffer. The supernatant was removed the following day, and the beads were washed once with 0.5 mL of isolation buffer. Then, 0.5 mL of plasma or EV fractions purified by SEC were added to the beads, which were incubated for one hour at 4°C with rotation in a Hula mixer. Afterward, the beads were washed twice with 0.5 mL of isolation buffer and then resuspended in 0.5% Triton X-100 in isolation buffer. This mixture was incubated for 10 minutes to release luminal proteins from EVs, and the supernatant was collected from the beads for further analysis.

## Data analysis

Data analysis was performed in Graphpad Prism 7. All figures were plotted in Graphpad Prism 7 and Adobe Illustrator version 2022.

## Competing Interest

David R. Walt is a founder and equity holder in Quanterix. His interests were reviewed and are managed by BWH and Partners HealthCare in accordance with their conflict of interest policies.

## Acknowledgements

The research described in this manuscript was supported by a grant from the Michael J. Fox Foundation (Grant number 2021A017224). Tal Gilboa is an awardee of the Weizmann Institute of Science Women’s Postdoctoral Career Development Award

