## Supporting Information for "Assessment of extracellular vesicle protein cargo as neurodegenerative disease biomarkers"

Table of contents:

Figure S1: Simoa α-synuclein assay
Figure S2: Measurement of α-synuclein in individual plasma samples and corresponding EVs isolated by SEC
Figure S3: Simoa pSer129 α-synuclein assay
Table S1: Spike and recovery α-synuclein and pSer129 α-synuclein assays
Table S2: Clinical characteristics of pooled plasma samples
Figure S4: Optimization of the protease protection assay
Figure S5: Protease protection assay for estimating the internal and external Tau in plasma EVs isolated by SEC

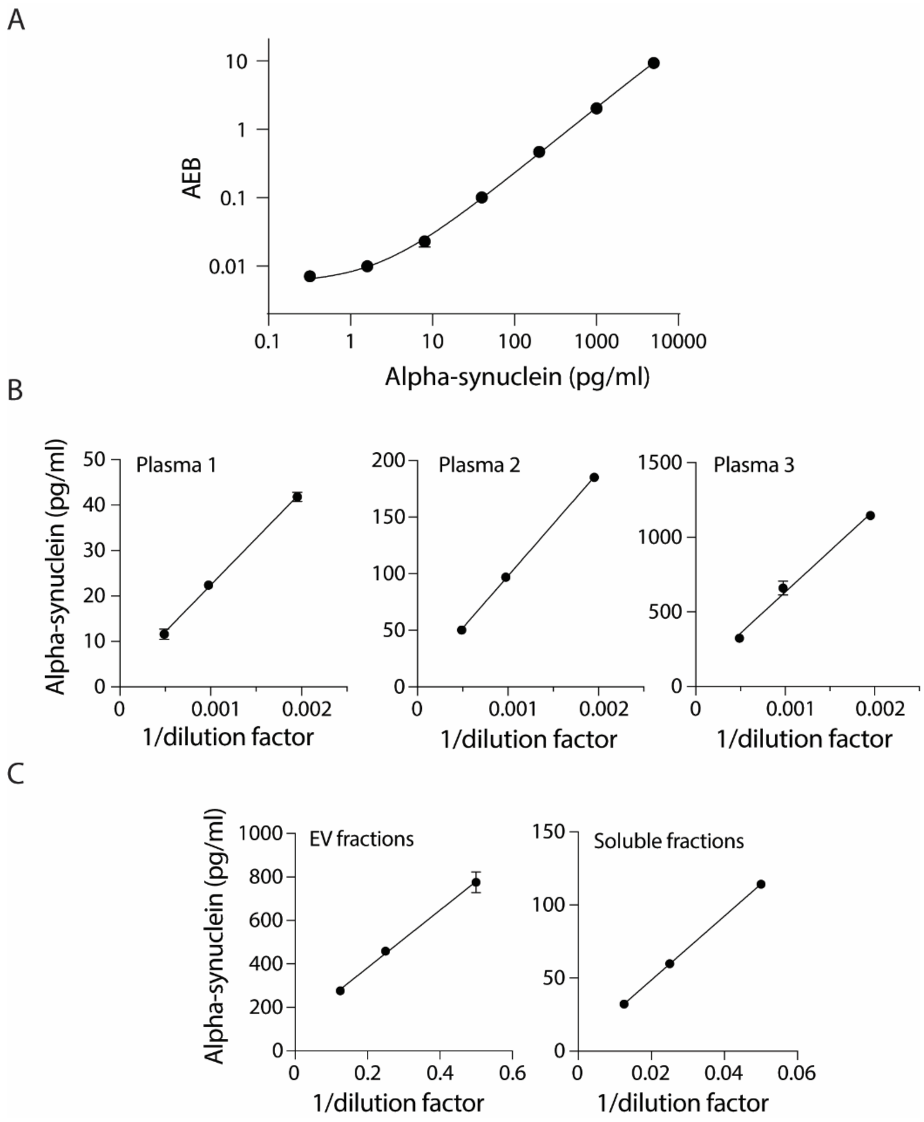


**Figure S1 | Simoa α-synuclein assay.** (A) Calibration curve of the α-synuclein Simoa assay. The recombinant protein standard was serially diluted. Error bars represent the standard deviation of duplicate measurements. A 4PL curve was fitted to the curve and the limit of detection (LOD) was estimated as 3 STD above background. LOD=0.41 pg/ml. (B) Dilution linearity was performed on three different pooled plasma samples. (C) The pooled plasma sample was fractionated and EVs fractions 7-10 (left), and soluble proteins fractions 17-20 (right) were collected and measured with three dilutions. All the data are displayed as mean ± SD.

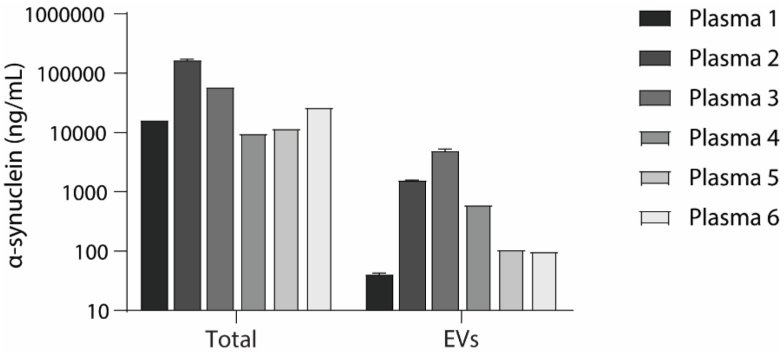


**Figure S2 | Measurement of α-synuclein in individual plasma samples and corresponding EVs isolated by SEC.** The level of α-synuclein was measured by Simoa in EVs isolated by SEC (fractions 7-10) from six individual plasma samples. Levels of each protein were also measured in corresponding unfractionated plasma samples. All the data are displayed as mean ± SD.

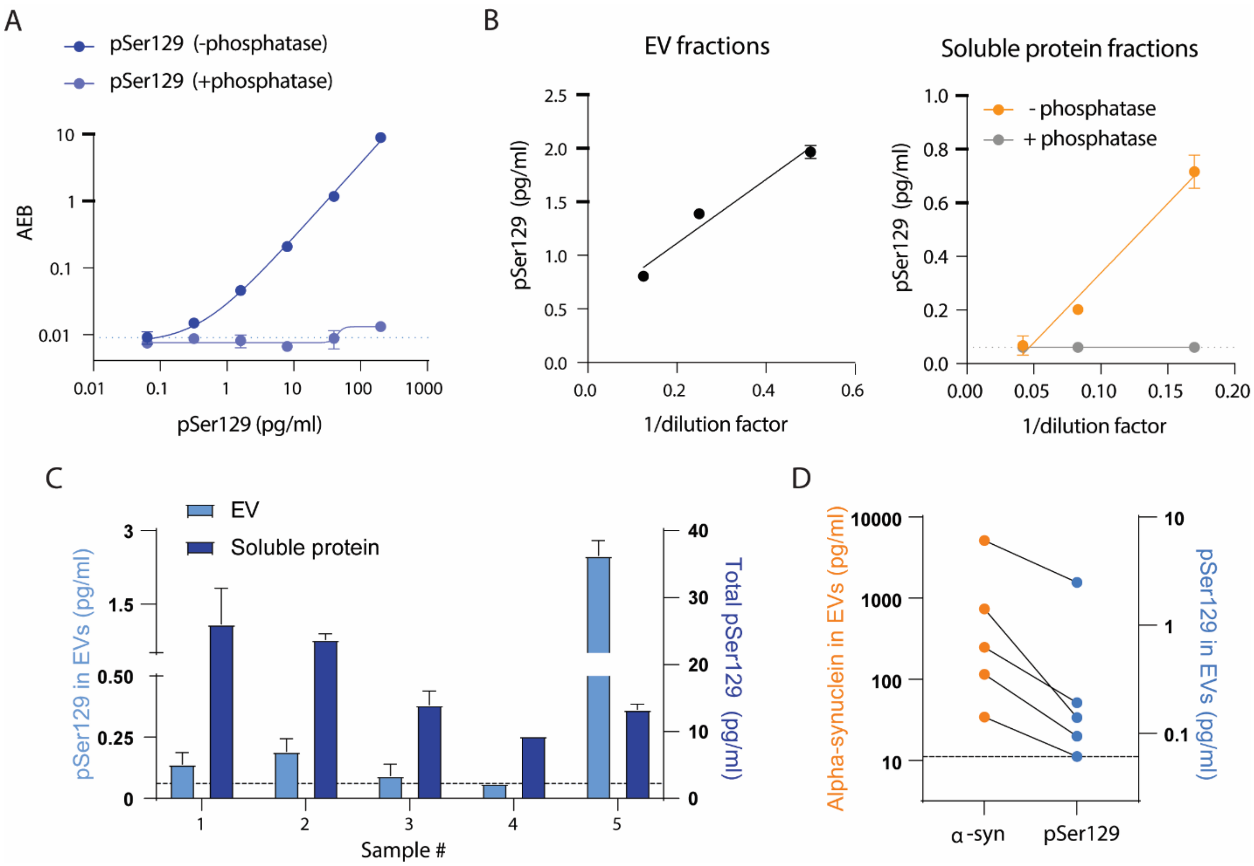


**Figure S3. Measurement of phosphorylated $\alpha$-synuclein at Ser129 residue (pSer129) in extracellular vesicle (EV) and soluble protein SEC fractions isolated from plasma.** (A) Purified pSer129 proteins were treated with (light blue) or without (dark blue) lambda phosphatase for 30 min at 30°C and then were titrated to calibration points. Calibration curve was performed with pSer129 Simoa assay. (B) Linearity of pSer129 assay was performed on pooled EVs (left) and pooled soluble proteins (right) isolated from four individual plasma by SEC. Mean linearity of EV fractions and soluble protein fractions is 128.6% and 61.8%. In order to confirm specificity of pSer129 assay, linearity was validated in presence of lambda phosphatase (gray dots), which showed signals below LOD (the dashed line). (C) Plasma of five healthy individuals was fractionated by SEC into EV and soluble protein fractions. The levels of pSer129 were measured in EV (light blue bars) and soluble protein (dark blue bars) fractions in Simoa. The levels of pSer129 were detectable in four out of five EV samples. LOD is displayed as the dashed line. (D) The levels of $\alpha$-synuclein (orange dots) were evaluated in the same cohort of EV samples as (C). Comparison of $\alpha$-synuclein and pSer129 is shown with lines connecting $\alpha$-synuclein signals to pSer129 signals in the corresponding EV samples. The dashed line shows LOD of pSer129 assay. All the data are displayed as mean ± SD.

| Simoa assay | Spike concentrations (pg/ml) | Recovery (%) |
|---|---|---|
| α-synuclein | 20 | 81.3 |
| | 100 | 80.9 |
| | 500 | 94.7 |
| pSer129 α-synuclein | 0.5 | 81.3 |
| | 2 | 108.0 |
| | 8 | 96.3 |

**Table S1 | Spike and recovery of α-synuclein and pSer129 α-synuclein assays in plasma.** Spike concentrations were chosen to be in the range of the concentration of the analyte in the plasma.

| Plasma Pool | N | Age (years) | Female (%) | Race | Ethnicity | Disease Duration (years) | UPDRS | MOCA |
|---|---|---|---|---|---|---|---|---|
| **Alzheimer's Disease** | 20 | 72.6 ± 11.0 | 40 | Black (10%), White (90%) | Not Hispanic (100%) | 6.4 ± 4.5 | N/A | 14.4 ± 7.2 |
| **Normal Control** | 20 | 74.8 ± 5.4 | 65 | Black (5%), Multiracial (5%), White (90%) | Hispanic (5%), Not Hispanic (95%) | N/A | N/A | 27.2 ± 2.1 |
| **Parkinson's Disease** | 20 | 72.5 ± 6.8 | 30 | Black (5%), White (95%) | Not Hispanic (100%) | 15.4 ± 5.7 | 34 ± 12 | 25.1 ± 4.5 |

**Table S2 | Clinical characteristics of pooled plasma samples**. Data expressed as mean ± standard deviation unless noted. N/A = Not applicable.

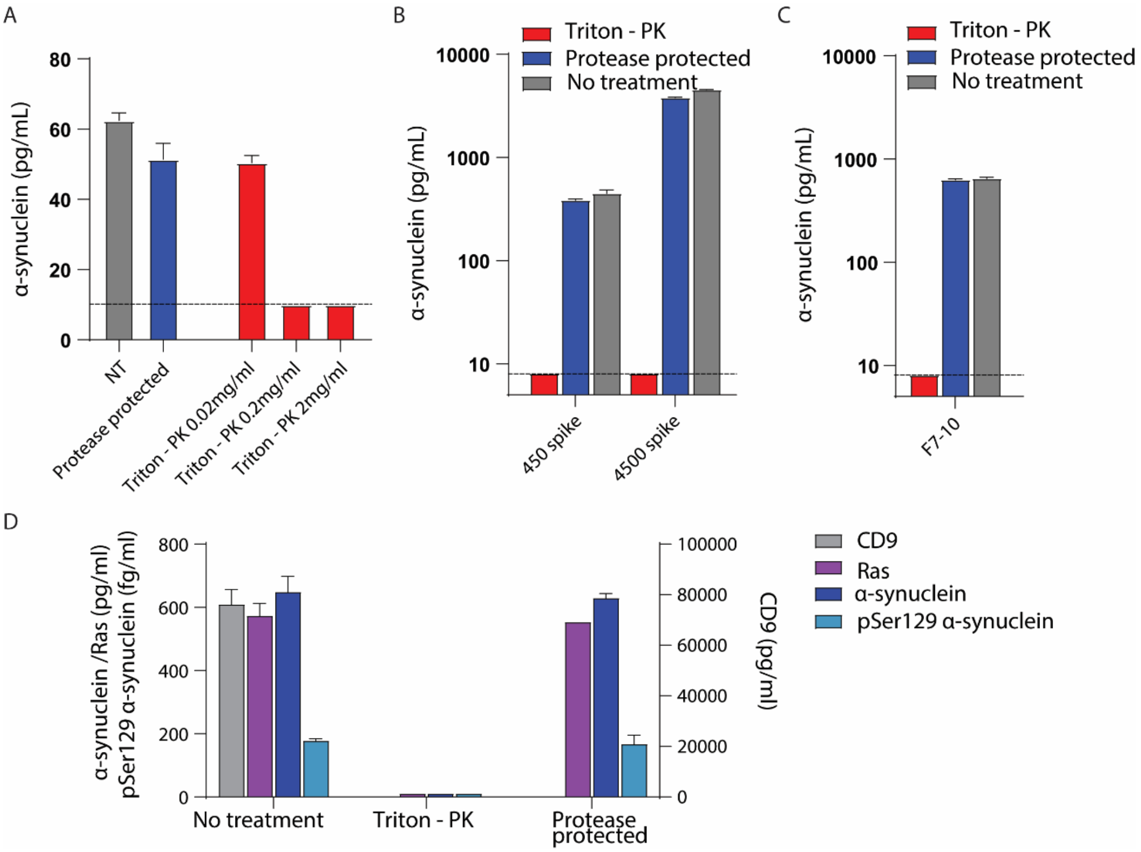


**Figure S4 | Protease protection assay optimization.** A) The proteinase K (PK) concentration was optimized on plasma EVs (SEC fractions 7-10) so that all the protein would be digested after lysing the EVs with Triton X-100. Simoa was then used to measure α-synuclein. B) Proteinase protection assay validation using two concentrations of α-synuclein protein standard (450 pg/ml and 4500 pg/ml). Protein standards were spiked into a PBS solution of inhibited PK (blue), active PK (red), or PBS only (gray). Simoa was then used to measure α-synuclein. (C) Protease protection assay validation using EVs isolated from plasma. EVs were isolated from pooled plasma using SEC (pooled fractions 7-10), and the sample was split into three conditions: no treatment (NT), proteinase protection assay, or PK treatment after EV lysis with Triton X-100 (Triton-PK). Simoa was then used to measure α-synuclein. The dotted line indicates the limit of detection for the α-synuclein assay. D) Validation of the protease protection assay using additional proteins. EVs were isolated from pooled plasma using SEC (pooled fractions 7-10), and the sample was split into three conditions: no treatment (NT), proteinase protection assay, or PK treatment after EV lysis with Triton X-100 (Triton-PK). Simoa was then used to measure α-synuclein, pSer129 α-synuclein, CD9, and Ras. As CD9 is an external marker, it was digested on both the Triton-PK and the protease-protected conditions. All the data are displayed as mean ± SD.

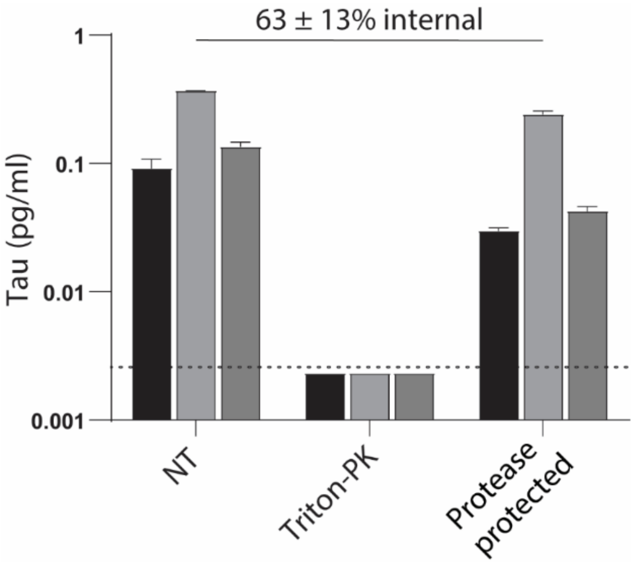


**Figure S5. Protease protection assay for estimating the internal and external Tau in plasma EVs isolated by SEC.** A protease protection assay was performed on pooled EV fractions (fractions 7-10) isolated from plasma. Levels of Tau in EVs isolated by SEC from three individual plasma were measured with Simoa. For each sample, three conditions were compared: No treatment (NT), protease protection assay, or PK treatment after EV lysis with Triton X-100 (Triton-PK). The dotted line indicates the limit of detection for the Tau Simoa assay. All the data are displayed as mean ± SD.
